# Eco-evolutionary feedback to a pesticide worsens the impact of a pesticide switch in a pivotal freshwater non-target species

**DOI:** 10.1101/2025.11.30.691396

**Authors:** Rafaela A. Almeida, Maxime Fajgenblat, Pieter Lemmens, Kiani Cuypers, Jade Maes, Kristien I. Brans, Luc De Meester

## Abstract

Pest management often requires the switches of insecticides, sequentially exposing non-target populations to different compounds. In a two-phased experiment, we assessed whether exposure to the pesticide chlorpyrifos induces rapid evolution in the non-target species *Daphnia magna,* and quantified the response of control and pre-exposed populations to a second exposure to the same pesticide, another pesticide with the same mode of action (malathion), or a pesticide with another mode of action (deltamethrin). Chlorpyrifos selection induced rapid shifts in genotype composition and reduced genotype richness, and strongly influenced population development following the second exposure. Chlorpyrifos-selected populations outperformed control populations when subsequently exposed to chlorpyrifos and malathion, but underperformed when exposed to deltamethrin. Our results highlight an eco-evolutionary feedback in which rapid adaptive responses to a pesticide worsens the response when exposed to a different type of pesticide in non-target species, increasing vulnerability to common agricultural practices.

## Introduction

Pesticides are strong drivers of adaptation in natural populations (Almeida et al., 2021; Hua et al., 2015; Siddique et al., 2020). Upon exposure to pesticides, populations can develop resistance through adaptive plastic and epigenetic responses (Brevik et al., 2018; Hua et al., 2013; Margus et al., 2019), allowing for rapid acclimation (DiGiacopo & Hua, 2020). Populations can also genetically adapt to better cope with pesticides (Almeida et al., 2021; Brans et al., 2021; Coors et al., 2009). Such genetic adaptations can happen through selection from standing genetic variation within populations, or through *de novo* mutations that confer higher resistance to pesticides (Hawkins et al., 2019).

While the capacity to adapt to pesticides has been well reported, adaptation to stressors can imply costs (Bourguet et al., 2004; Jansen et al., 2011d; van Kleunen & Fischer, 2005; Wang et al., 2010). Specifically, while adaptation to a pesticide is beneficial in its presence, such adaptation may lead to individuals underperforming when this stressor is no longer present (Jansen et al., 2011b; Lenormand et al., 1999) or leave adapted populations more susceptible to other stressors (Cuenca Cambronero et al., 2018; Jansen et al., 2011c). This increased susceptibility of stressor-adapted populations might be particularly important in the light of agricultural management schemes. Pesticide rotation is a common practice in pest management to counteract adaptation of a pest species to one particular pesticide and ensuring its effectiveness (Denholm & Devine, 2013). The frequent replacement of pesticides by newer compounds (Finger, 2018; Möhring et al., 2020) is also driven by the growing awareness of the adverse effects of specific pesticides (Butler, 2018; Zwetsloot et al., 2018), and by the increasing availability of new formulations (Bernhardt et al., 2017). Changes in pesticide also result from the growing popularity of organic farming, which is promoted as a more environmentally friendly alternative to conventional agriculture, but would also involve a shift in pesticide application, as the pesticides used in organic agriculture differ strongly from those in conventional agriculture (Gomiero et al., 2008). As a consequence, natural populations inhabiting agricultural areas are regularly exposed to new pesticides within one growing season. Negative effects of evolutionary adaptation to pesticides might in this context add to the burden of pesticide pollution on non-target species.

Pesticides impose strong threats to freshwater ecosystems (Peters et al., 2013; Rumschlag et al., 2020; Tang et al., 2021). Even pesticide concentrations below regulatory safety thresholds can impact freshwater populations (Liess et al., 2013; Liess & von der Ohe, 2005; Schäfer et al., 2007; Siddique et al., 2020). Farmland ponds are especially susceptible to pesticide contamination due to their location within agricultural areas and their small size. This is concerning, as pond ecosystems are collectively important biodiversity hotspots (Dudgeon et al., 2006) and provide key ecosystem services (Biggs et al., 2017). Within farmland pond communities, zooplankton can be strongly affected by pesticides (Hébert et al., 2021; López-Mancisidor et al., 2008; Relyea, 2005). In particular, species of the genus *Daphnia*, including *Daphnia magna*, have been widely used for decades as sentinel organisms in ecotoxicology and are embedded in standardized regulatory testing frameworks (OECD, 2004, 2012), and serve as a central model system in ecology and evolution (Ebert, 2005; Lampert & Kinne, 2011; Miner et al., 2012). Both lethal and sublethal effects have been reported for *Daphnia* in response to pesticide exposure, including reduced reproduction (Pestana et al., 2010; Song et al., 2017), delayed development (Toumi et al., 2013) and reduced grazing efficiency (Barata et al., 2007; Bengtsson et al., 2004). The latter finding is particularly important, as *Daphnia* play a central role in lentic ecosystems as key grazers, exerting strong top-down control on phytoplankton communities (Gianuca et al., 2016; Scheffer et al., 1993), thereby contributing to the prevention of toxic phytoplankton blooms. They also constitute a key prey resources for visual predators such as fish. Previous studies have documented adaptation to specific pesticides in *Daphnia* populations inhabiting farmland ponds (Almeida et al., 2021; Coors et al., 2009; Jansen et al., 2015; Simpson et al., 2015), underscoring their value for understanding long-term ecosystem responses to contamination.

Given that *Daphnia* populations can adapt to pesticide pollution, its potential costs in terms of susceptibility to other stressors, and the pesticide switches being a common agricultural practice, we designed an experiment to assess the effect of pesticide exposure on *Daphnia magna* populations that have been previously exposed to another or the same pesticide. We test the hypotheses that (i) exposure to a model organophosphate insecticide, chlorpyrifos, affects genotype composition of *Daphnia magna* populations, (ii) such exposure affects population dynamics when the populations are subsequently exposed to a new pesticide, and that (iii) this impact is dependent on a match between mode of action of the second to the first pesticide exposure. More specifically, we expect that selection by chlorpyrifos improves performance upon a second exposure to chlorpyrifos or exposure to malathion, another organophosphate pesticide with an identical mode of action. In contrast, selection through exposure to chlorpyrifos is expected to result in a lower performance in response to exposure to a pesticide with a different mode of action, such as deltamethrin. Following the recent EU ban on chlorpyrifos, pesticide replacements are expected in systems where this compound was previously applied. Deltamethrin is a pyrethroid pesticide with a different mode of action from organophosphates, and is chemically based on pyrethrins, which are used in organic agriculture. This switch scenario in the experiment was chosen to reflect a changing pesticide application strategy, as also seen in the current transition toward more organic agricultural practices.

## Material and Methods

### Clone isolation and rearing

We isolated 20 genetically unique clonal lineages of *Daphnia magna* by hatching resting eggs from the top sediment layer (upper 2 cm) of the pond Langerodevijver, located in a nature reserve in Flanders, Belgium (50°49’42.2” N – 4°38’23.7” E). These lineages were kept under standardized optimal laboratory conditions (20±1°C, 16:8 light:dark photoperiod, renewal of 75% of the medium and feeding with 1×10^5^ *Acutodesmus obliquus* cells/mL every second day) for at least two generations to avoid the interference from (grand)maternal effects.

### Pesticides

We selected three pesticides for this study: chlorpyrifos, malathion and deltamethrin. Chlorpyrifos (CAS 2921-88-2, purity > 99%, Sigma-Aldrich) is a broad-spectrum organophosphorus insecticide that has been commonly used in agriculture (Eaton et al., 2008; Racke, 1993) and acts as an acetylcholinesterase inhibitor (Solomon et al., 2014). Chlorpyrifos was banned by the European Commission in 2020 and has recently been listed for global elimination by the Stockholm Convention, hence pesticide switches stimulated by these regulatory bans are expected globally. Malathion (CAS number: 121-75-5, purity > 99%, Sigma-Aldrich), another organophosphate insecticide, and similarly to chlorpyrifos, is also an acetylcholinesterase inhibitor. Malathion has been extensively used in agriculture for the past half century (Jensen & Whatling, 2010), and even though its usage is restricted to greenhouses within the EU under the regulation (EC) No 1107/2009, it is still widely applied in crops and detected in waterbodies around the world (Vasseghian et al., 2021). Deltamethrin (CAS 52918-63-5, purity >98%, Sigma-Aldrich) is a synthetic pyrethroid insecticide (Soderlund, 2010) that acts on the voltage-gated sodium channels of nervous cells membranes (Field et al., 2017). Deltamethrin is widely used worldwide and, besides being allowed in organic agriculture in insect traps under the EU regulation (EC) No 889/2008 (Commission of the European Union, 2008), it shares the same mode of action and similar chemical structure with pyrethrins, the class of pesticides most commonly used in organic agriculture (Isman, 2006; Jansen et al., 2010). Based on pilot tests and earlier work (Almeida et al., 2021, 2023; Van de Maele et al., 2021), we chose environmentally relevant concentrations of chlorpyrifos, malathion and deltamethrin (0.35µg/L, 1.4µg/L and 0.075µg/L, respectively, Marino & Ronco, 2005; Mestres & Mestres, 1992; Vasseghian et al., 2022) that are expected to differentially affect *Daphnia magna* genotypes, inducing lethal effects on the most sensitive individuals as well as sublethal impacts on moderately tolerant individuals.

### Experimental design

The experiment was divided into two phases: a selection phase involving an exposure to chlorpyrifos for three weeks, followed by a second exposure phase in which an exposure to either no pesticide, chlorpyrifos, malathion, or deltamethrin was administered (Fig S1). The two phases of the experiment were separated by a ten-day recovery period to allow for density increase and reduce the impact of acclimatization.

#### Selection phase

In the selection phase, five juveniles (three to four days old) of each of the 20 clonal lineages were inoculated in 15L aquaria that were filled with 10L of dechlorinated tap water, resulting in six starting populations, with 100 individuals each. These populations were kept under standardized conditions (20°C, photoperiod 16:8 light:dark) and, after an acclimation period of three days, were exposed for three weeks to either a control treatment or a chlorpyrifos treatment (weekly pulses of 0.35 µg/L), each replicated three times (2 selection treatments x 3 replicates = 6 units). Populations were fed daily with 1×10^5^ *Acutodesmus obliquus* cells/mL. The aquaria were cleaned twice a week (on the day of a new pulse and three days after the pulse) by removing dead individuals and leftover algae to prevent deterioration of water quality. To ensure a weekly refreshed pulse of pesticide exposure, the total amount of medium was renewed once a week and a new pesticide pulse was given. After three weeks, a random 500mL sample was collected from each aquarium and preserved in absolute ethanol (>99.8%, Fisher Chemical) to isolate *Daphnia* for genetic analyses. After this, the populations were allowed a ten-day resting period, during which no pesticide was added to the aquaria and keeping the maintenance schedule identical to the experiment.

#### Second exposure phase

For the second exposure phase, the experimental populations from the selection phase were each divided into four populations by randomly isolating four sets of 70 sub-adult individuals from each aquarium. These populations were subsequently inoculated in 10L aquaria filled with 5L of dechlorinated tap water, and exposed to one of the four different treatments in triplicate: a control treatment (no pesticide exposure), exposure to chlorpyrifos (i.e. exposure to the same pesticide as during the selection phase), exposure to malathion (i.e. exposure to a pesticide with the same mode of action as chlorpyrifos) and exposure to deltamethrin (i.e. exposure to a pesticide with a different mode of action from chlorpyrifos). This design generated 2 selection treatments x 4 treatments in the second exposure phase x 3 replicates = 24 experimental units in the second phase. New pulses of each pesticide were given weekly (chlorpyrifos - 0.35µg/L, malathion — 1.4µg/L, and deltamethrin — 0.075µg/L), following the same procedure as during the selection phase, over three weeks. Laboratory conditions and feeding routine were identical to the selection phase. Population densities were determined twice a week by counting individuals based on video recordings of the *Daphnia* populations. The entire populations were transferred to shallow glass trays placed in a dark enclosure over a LED light pad (Huion L4S) with an overhead camera set-up (Canon EOS 700D). The individuals were recorded for ten seconds, and the video recordings were analyzed using the Trackdem R package (Bruijning et al., 2018; R Core Team, 2023). After three weeks, a random 500mL sample was collected from each aquarium and the animals were preserved in absolute ethanol for genetic analyses.

#### Genotyping

To genotype the individuals sampled from all experimental populations at the end of both the selection and the second exposure phase, we used eight microsatellite markers structured in one microsatellite multiplex (MO1) following Orsini et al. (2012) and Jansen et al. (2011a). Individuals were considered to be of identical genotype (i.e. belonging to the same clonal line) if they did not differ in their genotype across the eight studied microsatellite loci. Note that all clones we inoculated are genetically unique as they hatched from sexual (dormant) eggs. We performed genomic DNA extraction on 20 randomly picked adult *D. magna* individuals from each population, following a Proteinase K digestion method as described in Mergeay et al. (2008), followed by a qualitative PCR (QIAGEN multiplex PCR kit; QIAGEN, Netherlands; protocol detailed in Supplementary Information). Microsatellite alleles were scored with an ABI PRISM 3031 automated sequencer (Applied Biosystems) and analyzed with the Gene Mapper software (size standard Liz 500, Applied Biosystems).

### Data analysis

#### Genetic data

Given the clonal structure of *Daphnia* populations, we combined the allelic variant at every locus into a multi-locus genotype for each individual, and used these data to assess genotype richness and the relative abundance of the different genotypes in all populations. We used principal component analysis (PCA) to visualize differences in genotype composition among experimental populations.

Additionally, we used Bayes factors to formally quantify clonal selection throughout the experiment and clonal differentiation among treatments, acknowledging sampling uncertainty (see Supporting Information for methodological details). Following Kass and Raftery (1995), we interpret Bayes factors as follows: values between 1 and 3.2 indicate “evidence not worth more than a bare mention,” values between 3.2 and 10 “substantial evidence,” between 10 and 100 “strong evidence,” and values greater than 100 “decisive evidence”.

#### Bayesian hierarchical modelling of population growth curves

We used a hierarchical Gaussian process (HGP) regression approach (Rasmussen & Williams, 2006) to model population densities over time and to investigate the impact of an initial selection through exposure to chlorpyrifos and subsequent exposure to a particular pesticide on key population growth characteristics, including the maximal growth rate, average population density and maximum population density. Gaussian processes (GPs) are a non-parametric method to parsimoniously model time series (Rasmussen & Williams, 2006). They have been shown to strongly outperform conventional parametric growth models when dealing with nonstandard growth curves (Tonner et al., 2020).

Let *Y*_*u*,*t*_ be the population density of experimental unit *u* = 1, 2, … , 30 at day *t* = 0, 4, 7, 11, 14, 18, 21, with corresponding exposure during selection *p*(*u*) = control, chlorpyrifos and second exposure *e*(*u*) = control, chlorpyrifos, malathion, deltamethrin. We assume *Y*_*u*,*t*_ follows a negative binomial distribution:

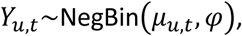

where *μ*_*u*,*t*_ is the expected density of experimental unit *u* at time *t*, and *φ* is a dispersion parameter common to all experimental units and time points. We model the linear predictor for each experimental unit as follows:

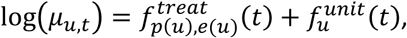

where 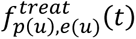 is a smooth function of time representing the actual growth curve for each the selection *p* and the second exposure *e* of experimental unit *u*, and where 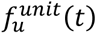 is a smooth function of time capturing the residual temporal pattern through time for each experimental unit *u*, not captured by the treatment-level function. Both groups of smooth functions are modelled by means of GPs:

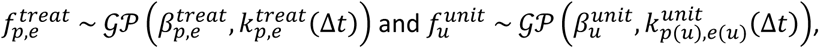

where 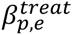 and 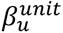 can be seen as treatment- and unit-level intercepts, and where 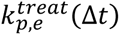 and 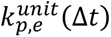 are covariance functions, dictating how fast similarity among any two pairs of measurements decays as a function of the time difference Δ*t* that separates them. We specifically consider an exponentiated covariance function, and we use weakly informative to informative priors on the model parameters (see Supporting Information for methodological details).

The HGP model allows us to probabilistically infer the true population growth trajectory for each combination of selection and second exposure condition, while simultaneously taking into account the temporally structured residual variation among replicates. We also quantified the maximum growth rate, the maximum population density and the average population density at each posterior draw to fully propagate uncertainty (see Supporting Information for methodological details), and we used a least squares approach to disentangle the effects of selection by chlorpyrifos exposure, second exposure to any of the three pesticides, as well as their interactions, across the eight treatment combinations.

We implemented the above model using the probabilistic programming language Stan and the rstan v.2.21.2 package (Stan Development Team, 2020) in R v.4.0.3 (R Development Core Team, 2020). Stan performs Bayesian inference by means of dynamic Hamiltonian Monte Carlo (HMC), a gradient-based Markov chain Monte Carlo (MCMC) sampler (Carpenter et al., 2017). We ran four chains with 2,000 iterations each, of which the first 1,000 were discarded as warm-up. We assessed model convergence visually by means of traceplots and numerically by means of effective sample sizes, divergent transitions and the Potential Scale Reduction Factor, for which all parameters had *R̂* < 1.01 (Vehtarh et al., 2021). We used the tidybayes v.2.3.1 package (Kay, 2020)(Kay, 2020) to visualize posterior distributions.

## Results

### Genetic changes following the selection experiment

Chlorpyrifos selection induced repeatable shifts in genotypic composition compared to the starting population and the control-selected populations (Figure 1, 2 and S2), with the first principal component of the PCA being able to clearly discriminate between genotype composition of control and chlorpyrifos-selected populations (Figure 2). Exposure to chlorpyrifos during the selection phase reduced genotype richness within populations (Figure S2), resulting in strong genetic differentiation between chlorpyrifos-selected populations and both the initial inoculation and control-selected populations at the end of the selection phase (Bayes factor 7.1×10^10^ and 6.9×10^9^, respectively, pooled replicates, Figure 1). Relative genotype frequencies in control populations also differed from those in the initial inoculation, although to a lesser extent (Bayes factor 4.6×10^10^, pooled replicates). More specifically, genotypes 2, 3, 8, 13 and 19 occurred at higher frequencies in the chlorpyrifos-selected populations than in the control populations, whereas genotypes 5, 9 and 16 occurred at higher frequencies in the control-selected populations (Figure 2, Figure S2). Genotype composition at the end of the second phase of the experiment shows strong differences (Figure 1 and S2), generally reflecting the differences that built up mostly during the selection phase. At the end of the second exposure phase, all populations initially selected with chlorpyrifos showed strong differences in genotype composition relative to control populations, regardless of the type of second exposure (Figure 1 and S3). Exposure to pesticides during the second phase led to strong differentiation between population that received the control treatment during the selection phase, with control-chlorpyrifos and control-deltamethrin populations differing the most (Bayes factor 3.6×10^4^, pooled replicates) (Figure 1). This genetic differentiation at the end of the second phase, however, was not observed in the populations with the chlorpyrifos selection.

**Figure 1.**
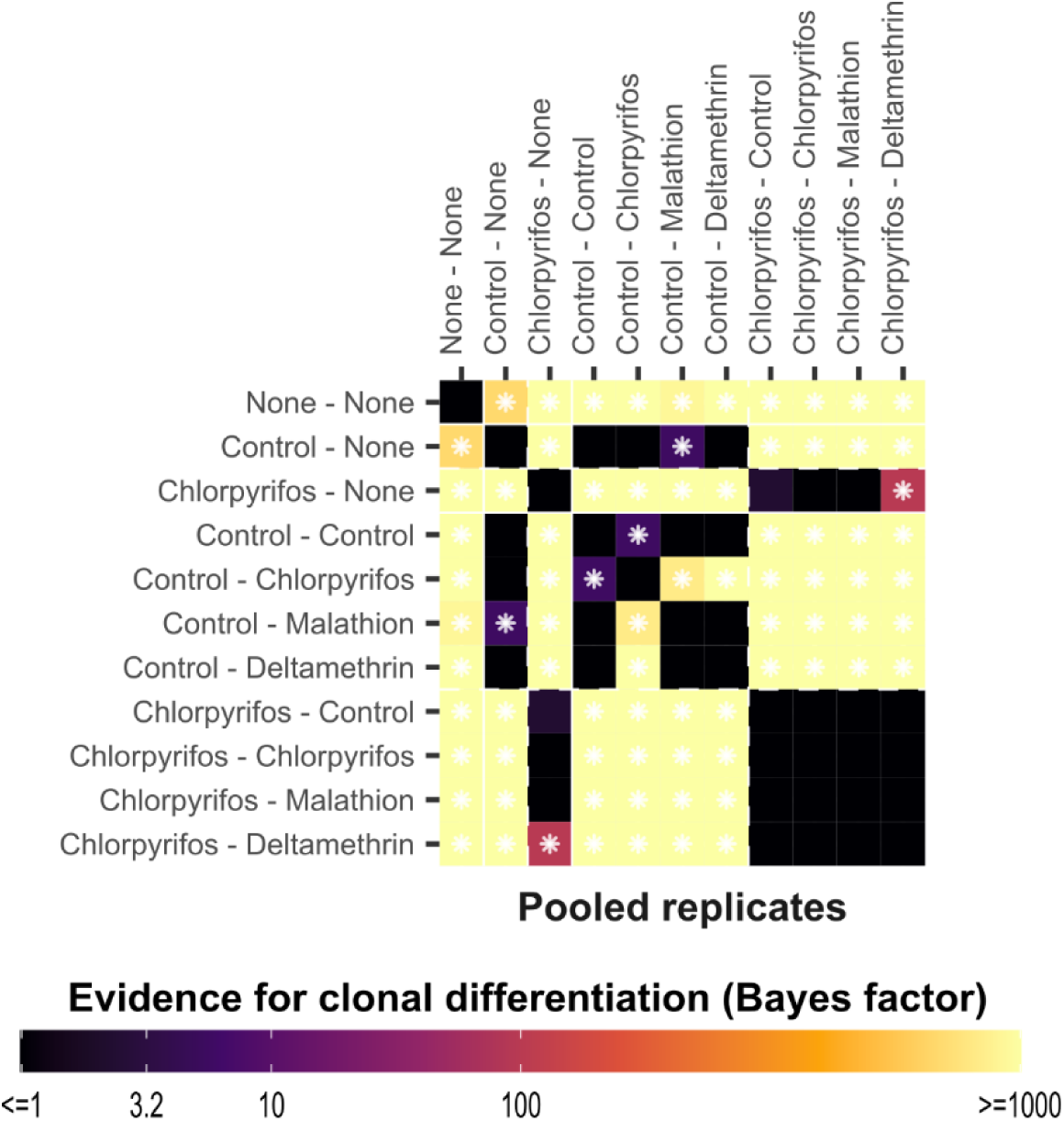
Heatmap showing the evidence for clonal differentiation among the genotypes of all pairs of experimental units in terms of Bayes factors, for the pooled replicates. Axis labels display the first and second exposure treatments, separated by a dash. The brighter the color, the more evidence for clonal differentiation. Pairs of experimental units that have at least substantial evidence for clonal differentiation (i.e. Bayes factor > 3.2) are highlighted by a white star.

**Figure 2.**
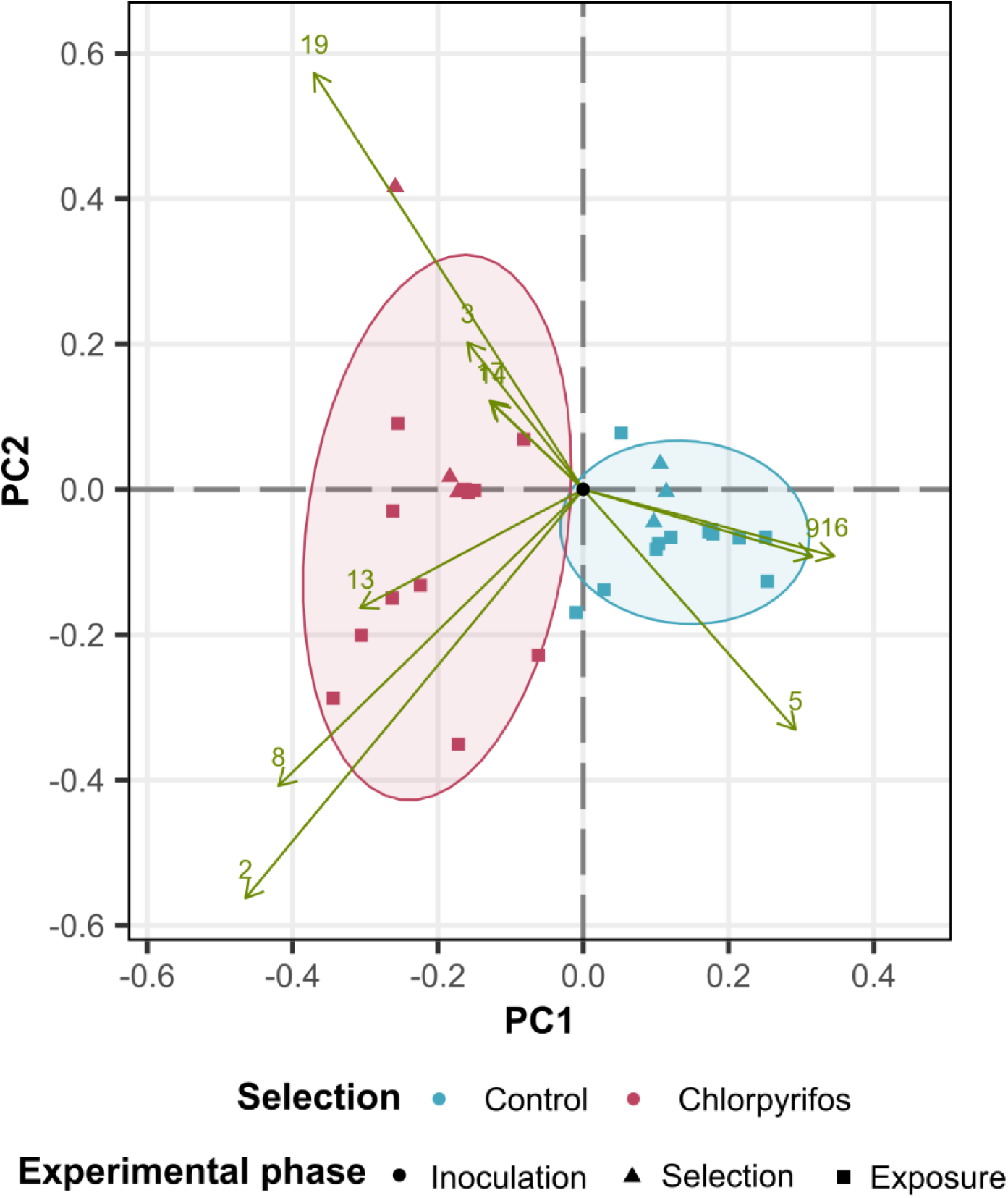
Principal component analysis (PCA) of the genotype composition of the populations of the different experimental populations among the two phases of the experiment. Biplot showing the ordination of populations along the first and second principal components, with colors depicting the selection phase type and symbols depicting the second exposure phase. Green arrows with numbered arrowheads depict the genotypes and their variable loadings. For clarity, only the 10 variables with the highest variable loadings are shown. The black point at the origin shows the ordination of the original community used for inoculation, with equal genotype frequencies.

### Population dynamics in the new pesticide phase

Our analysis revealed that chlorpyrifos selection strongly influences the response of the populations in the second exposure phase, especially the average and maximum population densities (Figure 3 and 4). Chlorpyrifos-selection increased maximum density during the second phase of the experiment by 650.8 individuals (posterior mean, 95% CrI [178.0; 1241.7], 99.4% posterior probability of an increase), average density by 199.5 individuals (95% CrI [9.3; 391.4], 97.8% post. prob. of an increase) and growth rate by 10.3 individuals/day (95% CrI [−4.0;35.7], 88.6% post. prob. of an increase) when compared to control-selected populations (Figure 3).

**Figure 3.**
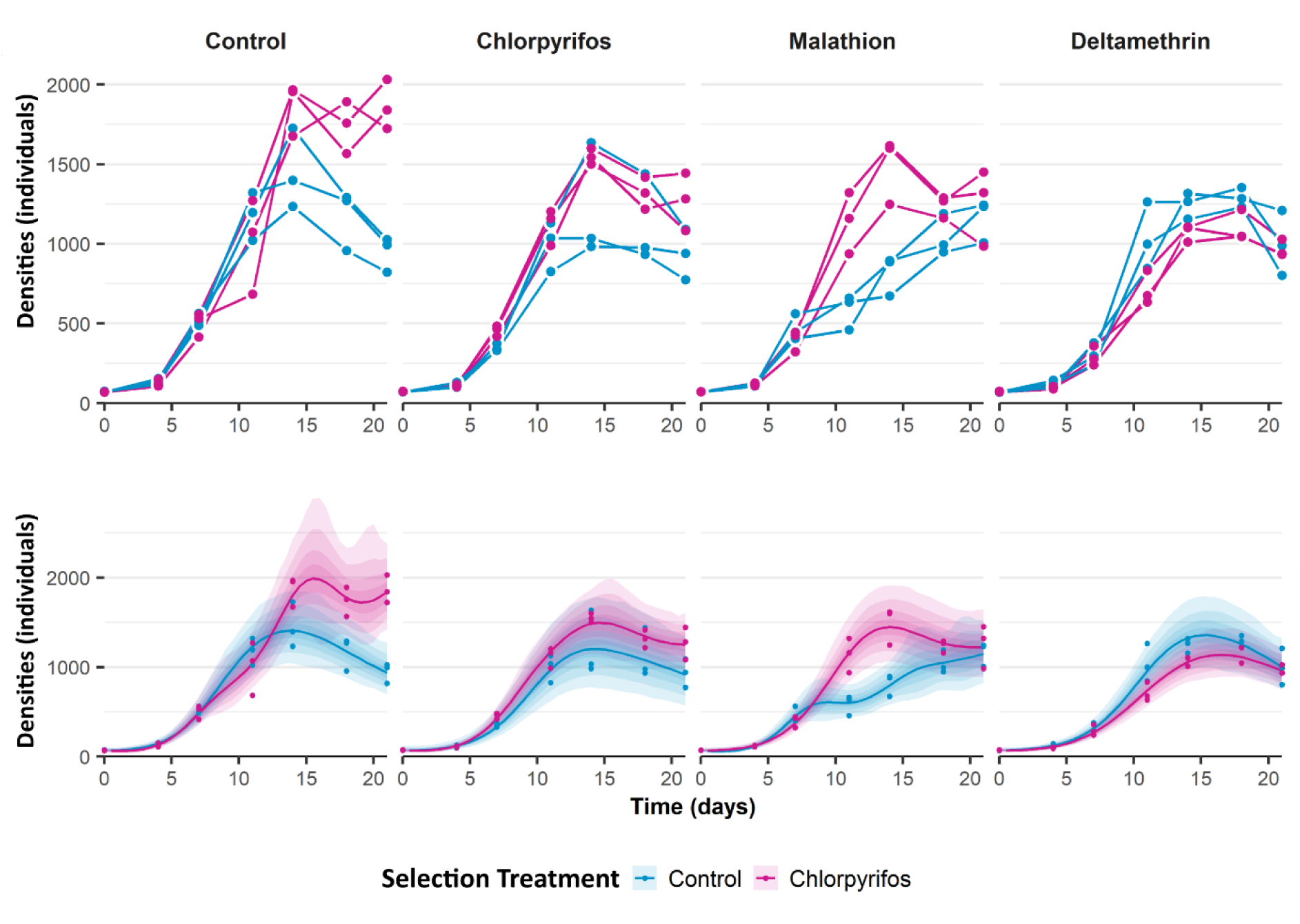
Observed (Top) and estimated (Bottom) population density patterns of Daphnia magna (number of individuals) that were either non-exposed (blue – control) or exposed to chlorpyrifos (pink) during the selection phase, throughout the second exposure phase of the experiment, when exposed to new control conditions, chlorpyrifos, malathion and deltamethrin. For the estimated densities, the full lines indicate the posterior median population densities, while the colored bands represent the 50, 80, 95 and 99% credible intervals. Original data are shown as points.

**Figure 4.**
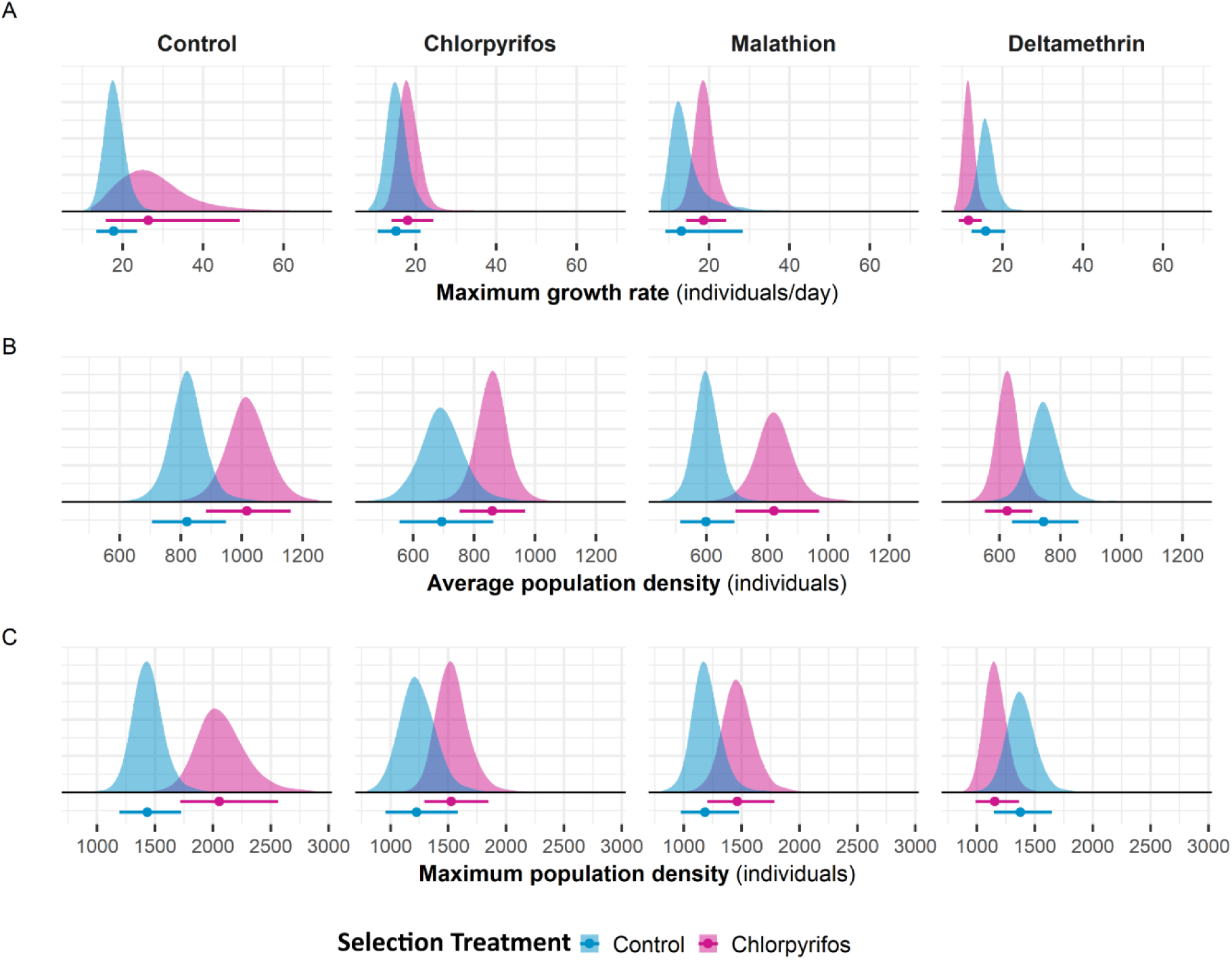
Posterior densities for the maximum growth rate, average population densities and maximum population density for each treatment during the second exposure phase of the experiment, for the populations that were non-exposed (blue – control) or exposed to chlorpyrifos (pink) during the selection phase. The horizontal bars represent 95% credible intervals, while the dots represent the posterior medians. Credible intervals quantify uncertainty around individual marginal estimates; treatment effects are instead evaluated from the posterior distributions of the treatment contrasts shown in Figure 5. The vertical scale corresponds to relative probability density and is omitted for clarity.

For populations exposed to control conditions in the second phase, previous chlorpyrifos selection increased maximum density by 46.6 percentage points (95% CrI [11.2;96.4]), average density by 25.0 percentage points (95% CrI [1.0; 53.2]) and growth rate by 60.5 percentage points (95% CrI [−20.4; 217.7]) when compared to the control-selected populations (Figure 4).

Exposure to any of the three pesticides (chlorpyrifos, malathion and deltamethrin) during the second phase tends to negatively affect populations, with consistently negative posterior median effects (Figure 5). Of the three parameters, the average population density is affected most: while there is a moderate signal for chlorpyrifos (−124.7 individuals, 95% CrI [−324.9; 77.2], 91.2% post. prob. of a decrease) and for deltamethrin (−75.0 individuals, 95% CrI [−232.5; 85.0], 85.4% post. prob. of a decrease), there is a clear signal for malathion in particular (−220.7 individuals, 95% CrI [−370.7; −63.9], 99.3% post. prob. of a decrease). We also identified a negative effect of malathion exposure on the maximum growth rate with moderately high probability (−3.3 individuals/day, 95% CrI [−11.5; 12.957], 81.8% post. prob. of a decrease).

**Figure 5.**
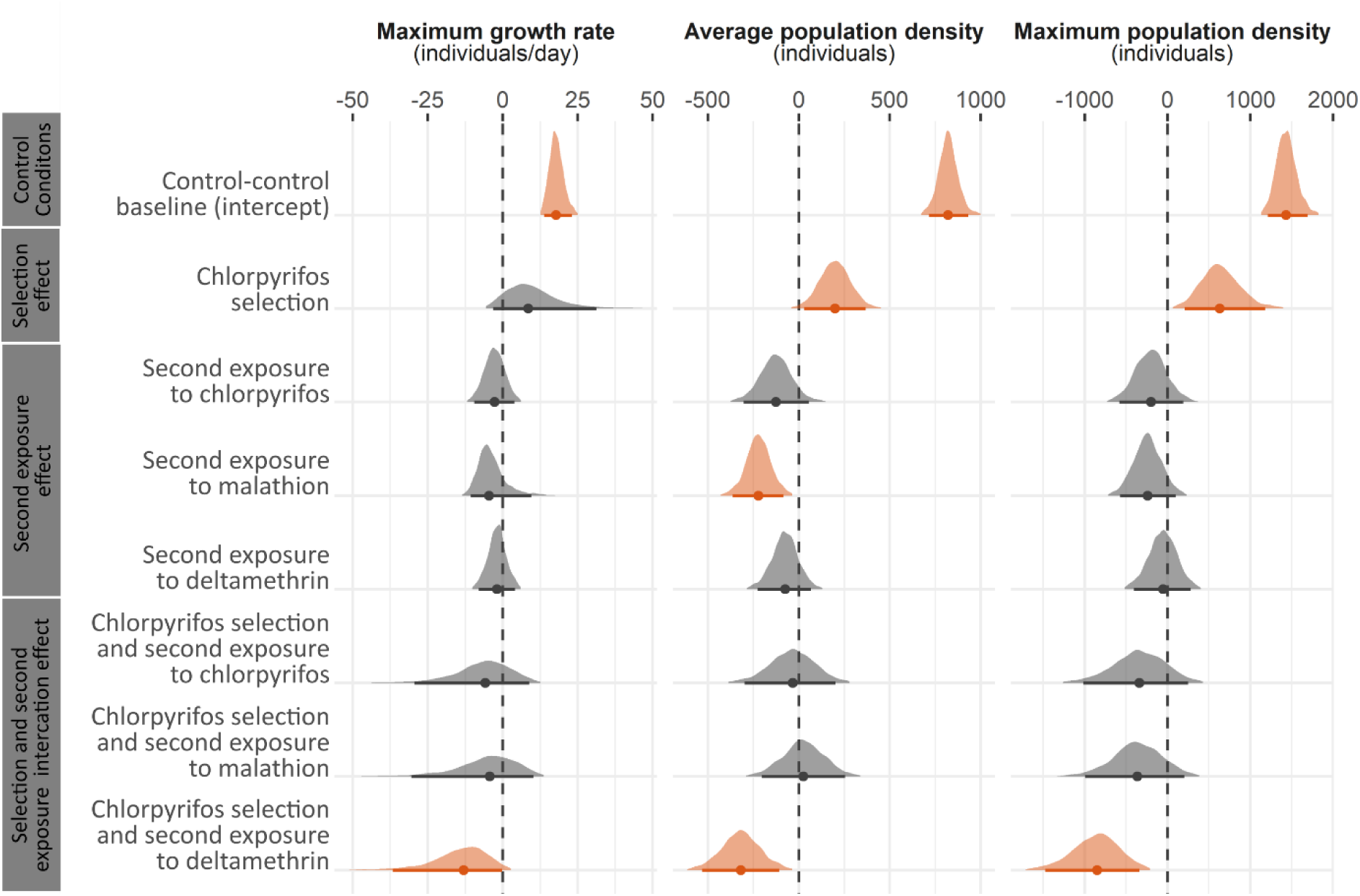
Posterior densities of the intercept of the control pre-exposure and control exposure baseline, the effect of chlorpyrifos selection, the effect of second exposure to chlorpyrifos, deltamethrin and malathion, as well as their interactions on the maximum growth rate, average population density and maximum population density of D. magna in the second phase of the experiment. Horizontal lines represent 95% credible intervals, and the points represent posterior medians. Posterior densities that have over 95% probability of being either smaller or larger than 0, are shown in orange. The displayed effects in this figure correspond to the quantities shown in Figure 4 as follows: each quantity in Figure 4 can be decomposed into the baseline intercept (corresponding to the control-control condition), the main effect of pre-exposure (if different from control), the appropriate main effect of exposure (if different from control) as well as the appropriate interaction effect if applicable. For instance, the total quantity for the chlorpyrifos-deltamethrin condition can be obtained by summing the intercept, the pre-exposure to chlorpyrifos effect, the exposure to deltamethrin effect as well as the interaction for chlorpyrifos selection and second exposure to deltamethrin.

There was a very strong negative interaction effect of chlorpyrifos selection followed by exposure to deltamethrin across all end points, namely growth rate (96.7% posterior probability), average density (99.5% posterior probability) and maximum density (99.8% posterior probability). More specifically, chlorpyrifos-selected *Daphnia* populations that were subsequently exposed to deltamethrin faced a 26.6 percentage point (95% CrI [−3.6; 49.6]), 15.7 percentage point (95% CrI [−1.6; 30.2]) and 15.5 percentage point (95% CrI [−7.3; 34.1]) reduction in maximum growth rate, average density and maximum density, respectively, compared to control-selected populations. In contrast, we did not find evidence for an important interaction of chlorpyrifos-selection followed by exposure to either chlorpyrifos or malathion.

## Discussion

Overall, exposure to chlorpyrifos during the selection phase strongly affected genotype composition and diversity within a three-week period, and subsequently affected demographic responses of the experimental *Daphnia* populations, both in the presence and absence of a second pesticide exposure. Whether chlorpyrifos selection had a positive or negative effect on population development upon a second exposure depended on the identity of the pesticide. In line with our expectations, population densities increased in chlorpyrifos-selected populations when exposed for a second time to the same pesticide or a pesticide with the same action mode (malathion), compared to control-selected populations. Population densities of chlorpyrifos-selected populations were negatively affected when exposed to deltamethrin in the second phase, compared to control-selected populations. Surprisingly, chlorpyrifos-selected populations also reached higher densities compared to control-selected populations when released from pesticide exposure in the second phase. Overall, our results indicate that *Daphnia* populations show rapid adaptive responses following exposure to a pesticide, and that this can lead to cross-tolerance but also to costs. This has important implications in the context of environmental policies that often lead to shifts in pesticide use.

A three-week exposure to the pesticide chlorpyrifos resulted in strong shifts in genotype composition in our experimental populations. Such shifts in clonal frequencies could result both from strong selection or from drift. However, the repeatable shift in genotype composition observed across replicates (Figure 1, 2 and S2) suggests a non-random response to selection. While we did not test tolerance of each genotype to chlorpyrifos, it is likely that exposure to chlorpyrifos lead to selection in favor of more tolerant genotypes or with higher tolerance plasticity. *Daphnia* populations can genetically adapt to pesticides (Bendis & Relyea, 2014; Jansen et al., 2011b, 2011c) as a result of *de novo* mutations (Chen et al., 2015; Gressel, 2011) or of selection on standing genetic variation within populations (Barrett & Schluter, 2008; Kersten et al., 2023), the latter allowing for faster responses (Hawkins et al., 2019). Given that we observe repeatable shifts in clonal frequencies, the response to selection in our experiment is mainly driven by standing genetic variation on tolerance or tolerance plasticity. As we used 20 genotypes hatched from dormant eggs of a single pond as the starting population, our experiment shows that the studied *Daphnia* population harbors substantial genetic variation relevant to the used selection factor. This is in line with the high capacity for (adaptive) evolution to another stressor (fish predation) that was found in a genomic resurrection ecology study (Chaturvedi et al., 2021). While our experimental design allowed for a ten-day period recovery from pesticide exposure and as such reduced impacts of direct pesticide exposure, we did not perform a strict common garden procedure involving multiple generations of purging from maternal effects in between the two phases of our experiment. As a result, we cannot without reservation claim that the differential responses of chlorpyrifos-selected *vs.* control-selected populations to a second pesticide exposure reflect exclusively the impact of evolutionary changes rather than acclimatization. Several studies have reported transgenerational epigenetic effects of pesticide exposure (Poulsen et al., 2021; Szabó et al., 2019), with exposed individuals and their offspring acquiring higher tolerance after exposure (Meng et al., 2025). While our observation of repeatable shifts in genotype frequencies linked to exposure to chlorpyrifos during the first phase of our experiment provides confidence that evolution likely played a role, we do not exclude that physiological acclimatization and epigenetic effects may have contributed to the observed responses in the second phase. For simplicity, we henceforth refer to the observed patterns, that might have resulted from both genetic as well as epigenetic effects, as adaptive responses. The responses are adaptive, as a first exposure to chlorpyrifos for three weeks resulted in a considerable improvement of the performance of the *Daphnia* populations when exposed to chlorpyrifos in the second phase of the experiment.

Similar to the pattern observed for chlorpyrifos, chlorpyrifos-selected populations performed better than control-selected populations when exposed to malathion in the second phase (Figure 3, 4 and 5). This suggests that beneficial traits for coping with chlorpyrifos might also increase population fitness under malathion exposure, reflecting cross-tolerance for these two organophosphates that act as acetylcholinesterase inhibitors (Jensen & Whatling, 2010). Cross-tolerance between pesticides of the same mode of action, and particularly between chlorpyrifos and malathion, has been reported previously across multiple organism groups (Bendis & Relyea, 2016; Hua et al., 2013; Saddiq et al., 2016; Van de Maele et al., 2021).

Surprisingly, chlorpyrifos-selected populations reached higher population densities under control conditions compared to those that were never exposed to a pesticide (Figure 3 and 4), with chlorpyrifos-selected populations reaching higher maximum and average densities in control conditions (Figure 4 and 5). These results contradict the findings that acquired resistance to a stressor reduces fitness under ancestral conditions (Kliot & Ghanim, 2012; Wang et al., 2010). One possible explanation for our observations might lie in changes in life history or physiological traits that might have been associated with the chlorpyrifos-induced shift in clonal composition. Studies on adaptive evolution of *Daphnia magna* to other stressors, such as heat, have shown effects on life-history traits. For example, urban-adapted populations evolved not only higher heat tolerance but also faster maturation, smaller body size, and higher intrinsic population growth rates than rural populations (Brans & De Meester, 2018). Furthermore, the observed response might also result from transgenerational plasticity and maternal effects, as pesticide exposure can affect reproduction of the non-exposed offspring generation (Castano-Sanz et al., 2022) and lead to smaller adults and offspring in *Daphnia* (Campos et al., 2016). Such genotypic or induced changes in life history strategy might also involve a differential investment of energy in eggs, as was observed by Glazier (1992) under food stress.

Populations that were first exposed to chlorpyrifos underperformed when exposed to deltamethrin (Figure 4 and 5). While we observed no apparent cost in the absence of a stressor, there seems to be a cost when exposed to a different type of pesticides. Several studies have shown that genetic adaptation to a given stressor can affect fitness when the population is exposed to a different stressor (Cuenca Cambronero et al., 2018; Dong et al., 2024; Jansen et al., 2011d, 2011c). Here, we observed a cost on all assessed demographic parameters: there was on average a 26.6 % decrease in maximal population growth rate, a 15.7% decrease in average and 15.5% decrease in maximal population density of populations that were previously exposed to chlorpyrifos compared to control populations.

Using a non-target species, we show that rapid adaptation to a pesticide reduces population performance in response to another pesticide. Applied to target species, this increases the effectiveness of pesticide switches, as it does not only reduce the likelihood of resistance evolution but also creates a cost when such adaptation already occurred. Applied to non-target species, however, this increases the harm done by pesticide use. Our results show that both increased resistance and associated costs in the response of *Daphnia* to pesticides may be observed within a very short time frame, namely a few weeks. Given that shifting pesticide use is a common agricultural practice, either to optimize pest management or in response to pesticide use legislation, the here reported eco-evolutionary feedback might be an important driver of negative impacts of pesticide use on natural populations and ecosystems.

In our study, initial selection under chlorpyrifos exposure followed by exposure to deltamethrin reduced genotype diversity and population densities of *Daphnia magna* populations. We did not observe complete population collapse nor a full recovery. The absence of a full recovery might reflect insufficient genetic variation or intrinsic trade-offs that limit adaptation. Some caution is needed to translate these results to real-world settings. First, our experimental set-up did not consider the complex community contexts in real landscapes and might hence have underestimated the impact of pesticide exposure on population performance (Almeida et al., 2023). On the other hand, our experiment started with a much lower genetic diversity than one expects in natural populations (Chaturvedi et al., 2021), so that the effect of genetic erosion might have played a larger role in our experiment than in nature. We note that genetic constraints will likely be much more important in many other organisms, such as threatened species that show reduced genetic diversity (Whitehead et al., 2017).

Our results highlight that frequent switches between different types of pesticides can impose substantial and previously underappreciated costs on non-target species. While cross-tolerance among pesticides has been described before (Bendis & Relyea, 2016; Hua et al., 2013; Van de Maele et al., 2021), our findings show that adaptive change in response to a given pesticide, associated with shifts in genotype composition, can translate into altered population-level demographic dynamics upon subsequent exposure to the same or another pesticide. We observed a striking cost in performance of populations that were previously exposed to chlorpyrifos and then to an insecticide with a different mode of action, deltamethrin. Our study therefore illustrates an important eco-evolutionary pathway through which exposure to pesticides might shape responses of non-target populations to changes in pesticide exposure, with the direction of this response being dependent on pesticide identity. This suggest an important evolutionary trade-offs, where adaptation to one stressor constrains responses to others (Fournier-Level et al., 2019; Kliot & Ghanim, 2012; Siddique et al., 2021). This eco-evolutionary feedback pathway is ecologically very important given the global application of pesticides in agriculture. Our results may be considered an example of what is likely a rather general observation: that the most effective strategies to suppress harmful species will likely also have the most negative effects on non-target species. Our results also indicate that, although ecosystems might benefit in the long-term from banning certain pesticides or transitioning to organic farming (Bengtsson et al., 2005; Nascimbene et al., 2012; Rundlöf et al., 2010), associated shifts in the type of pesticides applied can have short-term negative effects on non-target populations.

## Notes

### Competing Interest Statement

The authors have declared no competing interest.

### Summary of Updates

This version has a revised introduction and discussion

### Funding

- Fonds Wetenschappelijk Onderzoek (FWO), BE: 1228725N, 11E3222N

