## Supplementary information for "Eco-evolutionary feedback to a pesticide worsens the impact of a pesticide switch in a pivotal freshwater non-target species"

**Material and methods**

Hatching of resting eggs

The sediment was collected in April 2018. In the laboratory, the sediment collected from the pond was thinly laid in white trays filled with dechlorinated tap water, and kept at 20±1°C and a 16:8 light:dark photoperiod. The trays were checked daily for new hatchlings. These hatchlings were isolated and placed individually in separate 200mL glass jars filled with dechlorinated tap water to establish clonal lineages. These lineages were kept under the same standardized laboratory conditions: 20±1°C, 16:8 light:dark photoperiod, fed twice a week at a concentration of 1x10^5^ *Acutodesmus obliquus* cells/mL.

Pesticide preparation

At the beginning of the selection phase, we prepared a stock solution of chlorpyrifos dissolved in ethanol (CAS 64175, >99.8%, Fisher Chemical) at a concentration of 1mg/mL in a dark colored glass vial, which was stored in the dark at 4°C. For the second phase of the experiment, new stock solutions were made for chlorpyrifos, deltamethrin and malathion (ethanol used as a solvent). The final concentrations were prepared on the day on which they were added to the aquaria by adding the stock solutions to dechlorinated tap water.

Maintenance of experimental units

During both phases of the experiment, all aquaria were cleaned twice a week. Once a week, along with tank cleaning, the media were fully refreshed. For this, the population in each aquarium was sieved through a 0.64µm mesh filter, and the animals were then transferred to glass trays with dechlorinated 24h aged tap water. During the second exposure phase, these trays were used for video recording. Meanwhile, each tank was fully scrubbed to remove debris and leftover algae, cleaned with ethanol to remove any pesticide residue and rinsed with hot tap water at least three times before being refilled with new media. The experiment was carried out between September and December 2020.

Microsatellites analyses

Twenty *Daphnia* individuals were randomly picked from the samples collected at the end of each phase of the experiment and fixated in absolute ethanol (Fisher Scientific, CAS 64-17-5), and homogenized in 100 µl proteinase K-buffer (16mM [NH4]2SO4, 67 mM TrisHCl, pH 8.8, 0.01% Tween-20, 10% DTT and 0.5mM proteinase K). The samples were incubated overnight at 56°C, followed by a 10-min. denaturation at 96°C. We performed a qualitative PCR with the QIAGEN multiplex PCR kit (QIAGEN, Netherlands) using a T1 PCR machine (Biometra, Germany). Cycling conditions were the following: 95°C for 15minutes, 94°C for 30 seconds, 56°C for 1 minute and 30 seconds, 72°C for 1 minute and 30 seconds, 60°C for 30 minutes and 4°C to finalize, repeated for 27 cycli.

Bayes factor analysis to formally quantify evidence for clonal selection

We used a Bayes factor approach to formally quantify clonal selection throughout the experiment and clonal differentiation among treatments, acknowledging sampling uncertainty. For the experimental populations that underwent a first chlorpyrifos exposure, we compared the null hypothesis assuming identical genotype fractions (corresponding to the inoculum) against the alternative hypothesis assuming unequal genotype fractions reflecting clonal selection. We performed this standard Bayesian multinomial test on the observed genotype counts using the ‘multibridge’ v1.2.0 package (Sarafoglou et al., 2023a, 2023b) in R v4.3.1. We used a modified approach to quantify evidence for further clonal selection upon subsequent exposure, or to quantify genetic differentiation among any pair of experimental populations, since these comparisons involve two sets of observed genotype counts that both feature sampling variation. Specifically, for each pair of experimental populations, we compared the null hypothesis assuming that both populations were sampled from the same latent genotype distribution against the alternative hypothesis assuming they were sampled from different genotype distributions. In the null model, genotype counts in both samples are modeled as outcomes of a single multinomial distribution with a shared latent vector of genotype probabilities, while in the alternative model, the genotype counts are modeled as outcomes from two distinct multinomial distributions, each with its own latent vector of genotype probabilities. We implemented the null and alternative model in the probabilistic programming language Stan using the ‘rstan’ package v2.32.3 (Stan Development Team, 2023) and computed the Bayes factors using the ‘bridgesampling’ package v1.1-2 (Gronau et al., 2020) in R v.4.3.1 (R Core Team, 2023). In both analyses, we assumed a Dirichlet(1,1,…,1)-prior on the latent vectors of genotype probabilities. We computed Bayes factors for all pairs of experimental populations using pooled genotype counts among replicates, as well as for each replicate separately. The obtained Bayes factor values represent the ratio of the marginal likelihood of the alternative model to the null model. Following Kass and Raftery (1995), we interpret Bayes factors as follows: values between 1 and 3.2 indicate "evidence not worth more than a bare mention," values between 3.2 and 10 indicate "substantial evidence," between 10 and 100 "strong evidence," and values greater than 100 indicate "decisive evidence". Overall, this approach addresses several limitations of conventional Chi-squared tests to detect clonal differentiation: (1) it is robust to small and zero counts, (2) it allows for the direct comparison of two observed samples without relying on expected values, and (3) Bayes factors provide a direct measure of support for either the null or alternative hypotheses, avoiding the limitations of p-values (Wasserstein & Lazar, 2016).

Hierarchical Gaussian process regression: covariance functions and prior specification

We specifically consider an exponentiated quadratic covariance function:

$k_{p,e}^{treat}\left( \Delta t \right)={\alpha_{p,e}^{treat}}^{2}\exp\left( -\frac{1}{2}\left( \frac{\Delta t}{\rho_{p,e}^{treat}} \right)^{2} \right)$ and $k_{p,e}^{treat}\left( \Delta t \right)={\alpha_{p,e}^{treat}}^{2}\exp\left( -\frac{1}{2}\left( \frac{\Delta t}{\rho_{p,e}^{unit}} \right)^{2} \right)$,

where ${\alpha_{p,e}^{treat}}^{2}$ and ${\alpha_{p,e}^{unit}}^{2}$ are marginal variance parameters that control the amplitude in population density fluctuations on the log scale, and where $\rho_{p,e}^{treat}$ and $\rho_{p,e}^{unit}$ are length scale parameter that control the rate at which the covariance decays along with increasing ∆t.

We chose a weakly informative standard half-normal $\mathcal{N}^{+}\left( 0,1 \right)$ prior for the two sets of marginal standard deviation parameters $\alpha_{p,e}^{treat}$ and $\alpha_{p,e}^{unit}$ and an informative $\text{InvGamma}\left( 6.3, 1.9 \right)$ prior for the two set of length scale parameters $\rho_{p,e}^{treat}$ and $\rho_{p,e}^{unit}$, which places most prior mass on length scales between the shortest and longest distance between any two observed time points. We chose a weakly informative $\mathrm{StudentT}^{+}\left( 3,0,3 \right)$ prior for each of the treatment-level intercepts $\beta_{p,e}^{treat}$ and we model the experimental unit-level intercepts $\beta_{u}^{unit}$ by means of zero-centred random effects, with a weakly informative half-normal $\mathcal{N}^{+}\left( 0,1 \right)$ prior on the random effects’ standard deviation. Finally, we chose a vaguely informative $\mathrm{StudentT}^{+}\left( 3,0,10 \right)$ prior for the negative binomial distribution’s dispersion parameter $\varphi$.

Computation of derived growth function quantities

In addition to the seven time instances at which population densities were measured, we also inferred (i.e., interpolated) population densities for a regularly spaced grid of time points with a 12-hour interval throughout the duration of the study for visualization purposes and to compute three derived quantities at each posterior draw: the maximum growth rate, the maximum population density and the average population density. The maximum growth rate is calculated based on the finite difference method as the largest estimated increase in population density during a 12-hour interval. The maximum population density is calculated as the highest population density throughout the study period. The average population density is calculated as the average population density throughout the study period.

**Figures**

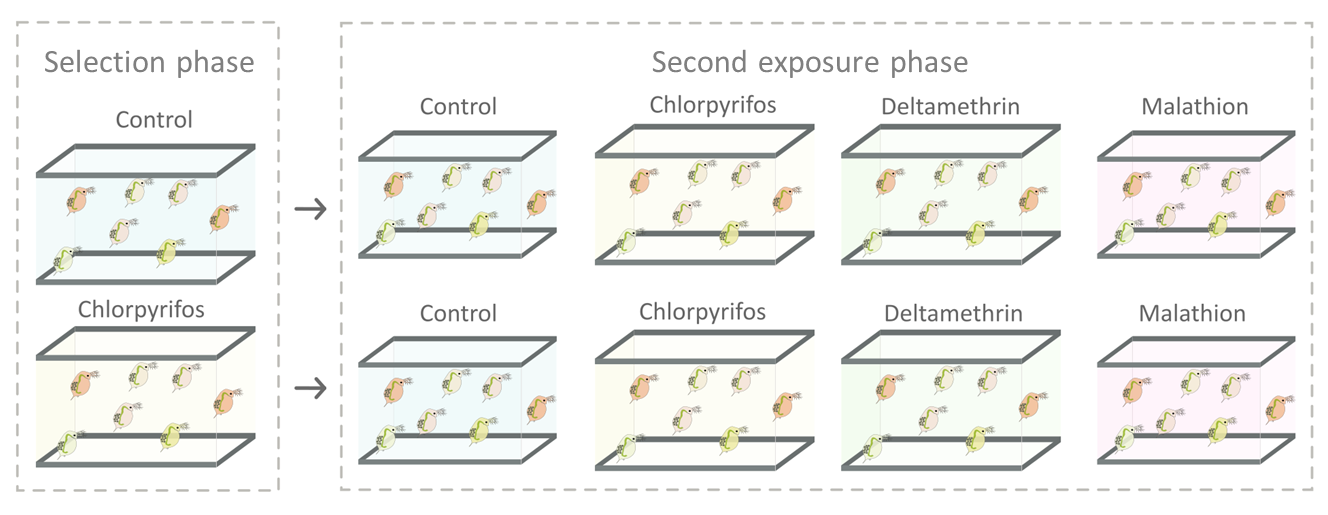

*Figure S1. Experimental design.* Daphnia magna *populations were first exposed to a selection phase, in which they were either kept in control conditions or exposed to chlorpyrifos, and subsequently exposed to a second pesticide exposure, in which the populations from the selection phase were split and exposed to one of the following conditions: control, chlorpyrifos, deltamethrin or malathion.*

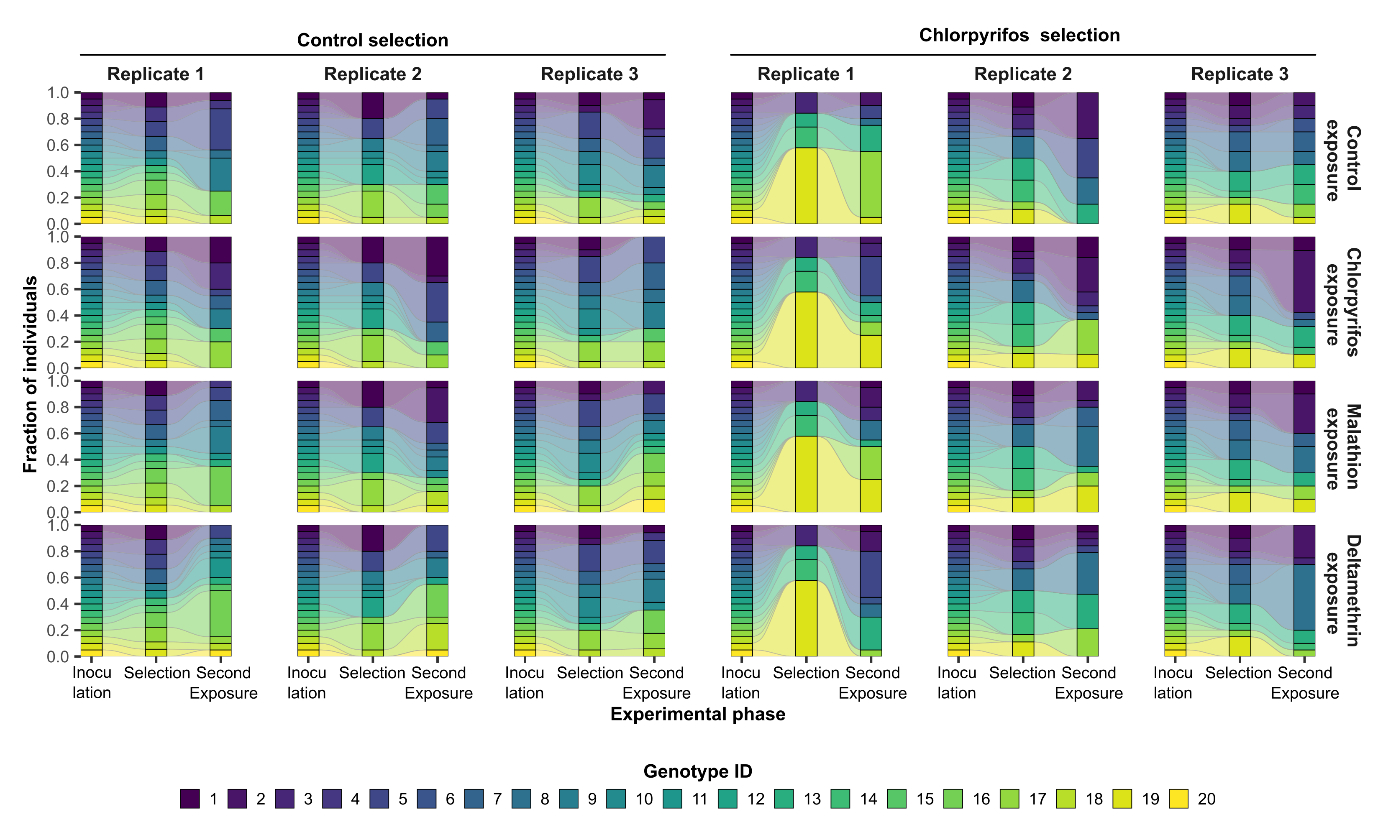

*Figure S2. Relative proportions of genotypes throughout the experiment, across treatments and replicates.*

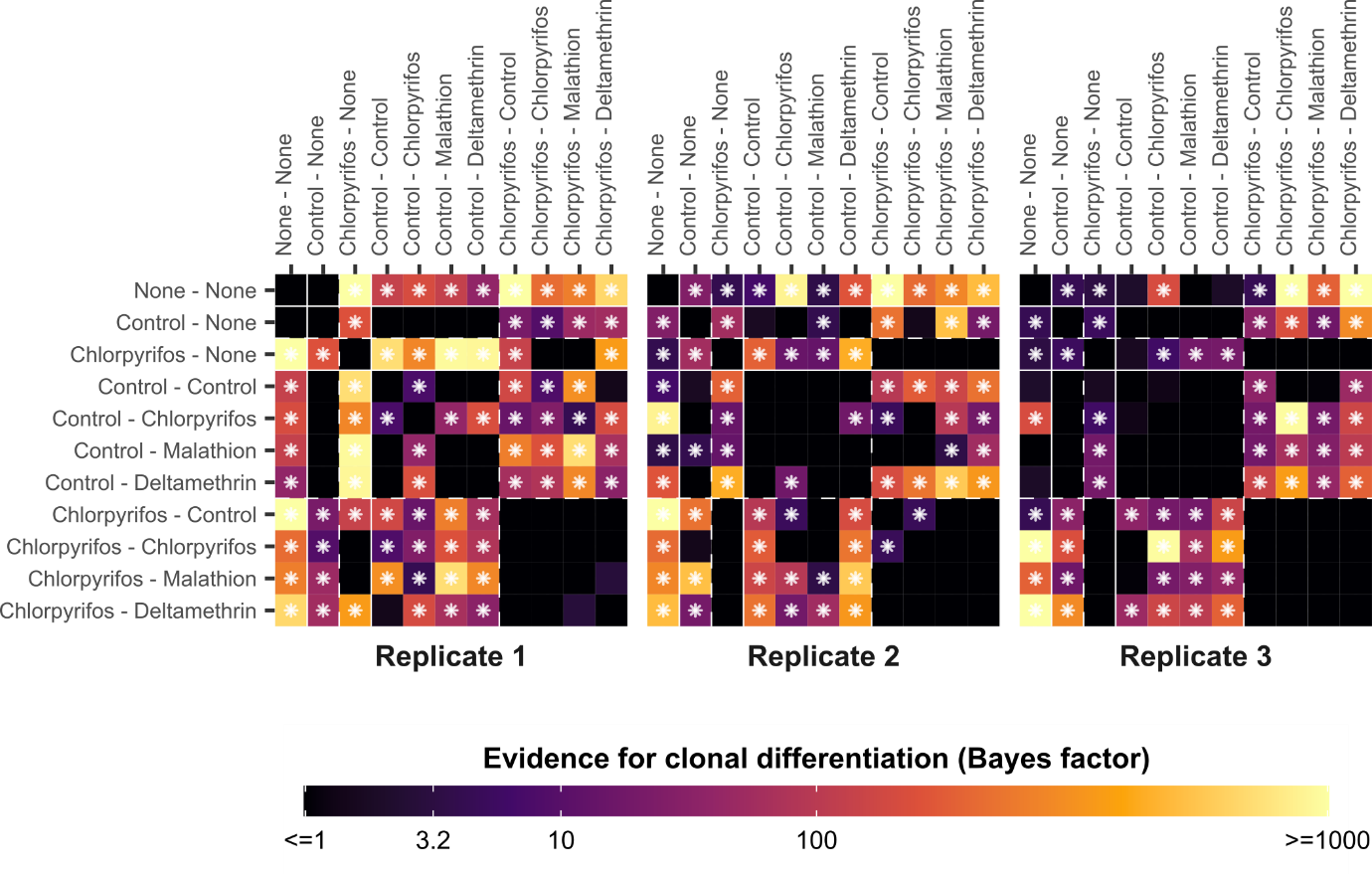

*Figure S3. Heatmap showing the evidence for clonal differentiation among the genotypes of all pairs of experimental units in terms of Bayes factors, for the individual replicates. Axis labels display the first and second exposure treatments, separated by a dash. The brighter the color, the more evidence for clonal differentiation. Pairs of experimental units that have at least substantial evidence for clonal differentiation (i.e. Bayes factor > 3.2) are highlighted by a white star.*
